# Time-separated Mutual Information Reveals Key Characteristics of Asymmetric Leader-Follower Interactions in Golden Shiners

**DOI:** 10.1101/2024.03.05.583541

**Authors:** Katherine Daftari, Michael L. Mayo, Bertrand H. Lemasson, James M. Biedenbach, Kevin R. Pilkiewicz

## Abstract

Leader-follower modalities and other asymmetric interactions that drive the collective motion of organisms are often quantified using information theory metrics like transfer or causation entropy. These metrics are difficult to accurately evaluate without a much larger amount of data than is typically available from a time series of animal trajectories collected in the field or from experiments. In this paper, we use a generalized leader-follower model to argue that the time-separated mutual information between two organism positions is a superior metric for capturing asymmetric correlations, because it is much less data intensive and is more accurately estimated by popular *k*-nearest neighbor algorithms than is transfer entropy. Our model predicts a local maximum of this mutual information at a time separation value corresponding to the fundamental reaction timescale of the follower organism. We confirm this prediction by analyzing time series trajectories recorded for a pair of golden shiner fish circling an annular tank.

## 1. Introduction

For many groups of social organisms, collective motion tends to be precipitated by one or more “leaders” who act first in response to a stimulus (sensing a food source, detecting an encroaching predator, etc.) and seemingly drive the rest of the group to follow their example. Seemingly is the proper qualifier, because although numerous studies have attempted to quantify the collective benefits of leader-follower dynamics [1–4] and identify behavioral features that make certain organisms more likely to take the lead in different scenarios [5–8], it is rather difficult to prove that the organisms who act later are actually being influenced by the leader and are not merely making independent, albeit delayed, decisions [9, 10]. Indeed, one can think of leader-follower behavior as a continuum, with organisms that blindly follow the leader at one extreme, and those that make independent decisions simply coinciding with the leader’s at the other.

To illustrate this latter extreme, consider a group of runners competing in a race. At any moment after the starting gun, one individual will be in the lead, and the other competitors will apparently be following this leader; but the reality, of course, is that each runner moves independently toward the finish line while jockeying for lead position. In this example, we understand that a true leader-follower modality is not playing out based on our contextual knowledge of the situation, but how is one to make a similar determination from observations of a flock of ducks flying in their characteristic v-formation or a mass of ants marching in single file?

At minimum, a leader-follower interaction must be asymmetric and retarded in time: the leader must influence the motion of the follower more strongly than vice versa, and the follower cannot react instantaneously to the motion of the leader. These requirements suggest that a necessary (but not sufficient) condition for the existence of a leader-follower relationship is a disparity in the size of the correlations between the present motion of one individual with the past motion of the other. The correlations should be large when comparing the present motion of the putative follower with the past motion of the presumed leader and should be small (ideally near zero) in the opposite case.

Currently, one of the most prominent metrics used to quantify these types of correlations is the transfer entropy [11–14]. For a pair of discrete-time Markov chains *X* and *Y*, the transfer entropy from process *Y* to process *X* is defined as [15]:

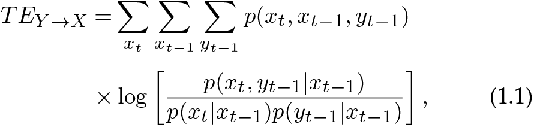

where *x*_*t*_ is the value of the Markov process *X* at time *t*, denoted *X*_*t*_, and *p*(*x*_*t*_) is its probability distribution function. This transfer entropy measures the amount of information shared between *X*_*t*_ and *Y*_*t−*1_, given that *X*_*t−*1_ is known. If we understand the conditioned random variable *X*_*t*_|*X*_*t−*1_ as encoding the stochastic dynamics of the process *X*, i.e., how the process chooses its value at time *t*, given its previous value, then *T E*_*Y →X*_ quantifies how much the dynamics of *X* are informed by the past state of *Y* .

Though popular, the transfer entropy has a number of shortcomings that make it difficult to accurately compute from experimental datasets. Most obviously is the fact that the trajectories of living organisms are not discrete-time Markov chains. While there are ways of generalizing the notion of transfer entropy to continuous time systems with a finite memory [16, 17], conditioning over a larger number of past states necessitates numerically estimating a higher-dimensional joint distribution (see Eq. (1.1)). This is less of an issue for an *in silico* model that can be simulated for arbitrarily long periods of time to produce datasets that representatively sample this high-dimensional multivariate distribution; but for experimental measurements of animal motion that often consist of a limited number of short time series, it is a serious problem [18].

One common solution is to determine a mesoscopic time scale *Δt* over which the dynamics of the experimental system are approximately Markovian and then compute the transfer entropy by replacing *t −* 1 in Eq.(1.1) with *t − Δt*. Again, however, data scarcity tends to preclude a rigorous estimation of this time interval, so oftentimes one settles for selecting a time scale that is known to be fundamental to the motion of the organisms under consideration [11]. The hope is that this physically motivated time scale will be sufficient to capture any asymmetry in the time-delayed influence between two organisms to determine whether a leader-follower type interaction between them is likely to exist.

Because of the higher-dimensional joint probabilities involved, accurately estimating the transfer entropy is a nontrivial task. The *k*-nearest neighbors estimator for mutual information first proposed by Kraskov, Stögbauer, and Grassberger [19] (henceforth abbreviated as the KSG estimator) has become the standard in many information theory studies [20–22]; but, although the algorithm can be straightforwardly generalized to estimate transfer entropy, it has been shown that this generalization reduces the accuracy of the method [23]. Alternative estimation schemes do exist, such as the use of kernel density estimators, but comparative studies have found them to perform, at best, only marginally better than the KSG method [24, 25].

In this paper, we argue for the use of time-separated mutual information as a viable alternative to transfer entropy for the purposes of identifying leader-follower interactions between pairs of organisms. For the Markov processes *X* and *Y* defined previously, this mutual information is defined as follows:

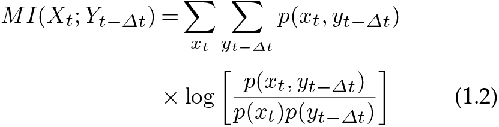

Though Kaiser and Schreiber [16] used a simple Boolean **3**stochastic process to contend that time-separated mutual information lacks the requisite asymmetry needed to correctly characterize leader-follower modalities (in their model, *MI*(*X*_*t*_; *Y*_*t−*1_) = *MI*(*Y*_*t*_; *X*_*t−*1_)); but we demonstrate that their finding was pathological and does not hold for a general class of leader-follower model that better represents the kinds of systems that are of interest in the study of collective motion. In comparing the motion of one organism at time *t* and another at *t − Δt*, we also avoid the problem common to transfer entropy studies of having to somehow choose the time scale *Δt*. Instead, we treat *Δt* as a variable and prove that, for pairs of organisms whose motion is adequately well described by our generalized modeling framework, the mutual information will peak at some critical value *Δt ≡ T*, which we identify as a measure of how quickly the follower organism can respond to a change in the motion of the leader, i.e., the organism’s reaction time.

We then test the predictions of our modeling framework by analyzing the motion of pairs of golden shiners (*Notemigonus crysoleucas*) swimming around an annular tank. We first discuss the formal requirements for correctly computing statistical metrics from a single time series and how laboratory experimental conditions can be chosen to fulfill those requirements. We then describe the specifics of our fish experiments and the results of our statistical analysis. For a well-behaved trajectory in which the two fish swam smooth, staggered laps around their tank, we use the time-separated mutual information between the angular positions of the two fish to demonstrate that their fluctuating angular separation is consistent with a genuine leader-follower dynamic with a well-characterized interaction time scale.

## 2. Methods

### (a) Generalized Leader-follower Model

In this section, we wish to encode our understanding of what a leader-follower interaction should minimally entail into a simple yet generalizable modeling framework. To that end, we consider two agents–a leader and a follower–whose configurations are, respectively, described by the vectors of random variables 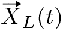 and 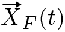. For point particle agents, their configuration is simply their vector position; for more realistic agents, the configuration may also include elements such as orientational coordinates, deformation coordinates, etc. The key point is that these two vectors evolve in discrete time according to the following pair of iterative equations:

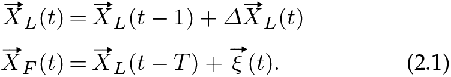

In the above, all times are given in units of an arbitrary (presumably small) time step and 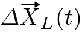 is a random variable representing the change in the configurational variables 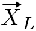 between consecutive time steps *t* and *t −* 1.All of the dynamics driving the motion decisions of the leader organism are modeled through 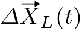, and we assume nothing about the distribution of this random variable other than that it does not depend explicitly upon the configuration of the follower.

The follower, on the other hand, attempts to pursue the same course as the leader, but its finite reaction time results in it lagging behind by *T* time steps. The follower cannot tail the leader perfectly, however, and its tendency to err is quantified by the random vector 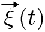. Again, we assume very little about the distribution of this noise term–only that its mean and variance must both be small on the scale of the domain through which the agents are moving.

We now consider the time-separated mutual information 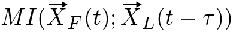 as a function of the time separation *τ*. We can relate 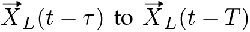 by iterating the first line of Eq. (2.1) either forwards in time or backwards, depending on whether *τ* is later or earlier than time *T*. The second line of Eq. (2.1) can then be used to relate 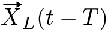to 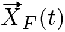as follows:

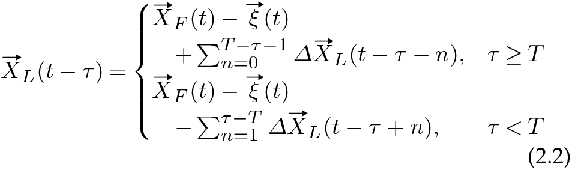

The above equation enables us to express 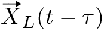as 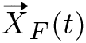 plus or minus a sum of terms that, by construction, do not depend upon the configurational state of the follower. This implies that as |*T − τ* | increases, the time-separated mutual information must decrease, since each additional term of the form 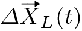simply contributes additional information not shared with 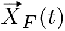. (Note that this is true regardless of whether these random variables are added or subtracted, though the actual quantitative value of the mutual information will typically be different in each case.) This means that as a function of *τ*, the time-separated mutual information for models of this class should have a peak value at *τ* = *T* and then decay towards zero as *τ* either increases or decreases away from *T* .

This parameter *T* physically represents an effective signaling timescale quantifying how long it takes information about the motion of the leader to travel to the follower, get processed cognitively, and then produce an observable response. For situations where the transmission of information and cognitive processing can be assumed to be of comparatively negligible duration (such as in the case of visual communication, wherein the information propagates at the speed of light), the quantity *T* can be understood simply as the reaction time of the follower.

If the model is at steady state, then we can exploit time translational invariance to show that the index-swapped mutual information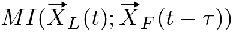 can be obtained by reflecting 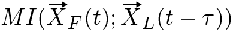 across the ordinate axis, i.e., replacing *τ* with *−τ*, and then translating both time arguments by *τ*. Since the time-separated mutual information is not (except in very special cases) a symmetric function about *τ =* 0, it is evident that 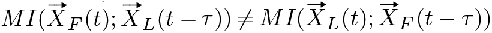, and the inherent asymmetry of the leader-follower dynamics is adequately captured by this two-point correlative metric.

### (b) Generating Statistics from a Single Time Series

Before we discuss the experimental system used to validate our generalized leader-follower model, it is worth discussing some of the difficulties inherent to computing information theory metrics from experimentally generated trajectory data and how designing experiments with these concerns in mind can help mitigate their impact. For a model system that is simulated *in silico*, it is generally not an issue to produce a large volume of replicate trajectories, each initiated from the same starting distribution of system configurations. For any time *t* after the start of the simulations, one can treat the corresponding time points of each replicate trajectory as statistically independent samples of the underlying configurational distribution at that time point. So long as the number of replicates is large enough to sample this distribution representatively, any statistic of the distribution can be computed accurately.

For trajectory data generated from the recorded motion of real organisms, producing such a high volume of replicates is infeasible, and so one often has no alternative but to treat points taken from the same time series as independent samples. This is only a reasonable assumption under two conditions. First, the system must be at steady state, so that the underlying distribution of configurations is stationary in time. Second, the time points selected as samples must be chosen far enough apart in time that any correlations between them have decayed to zero.

Many stochastic processes are inherently nonstationary, even at long times, but one can often force these systems to settle into a longtime steady state by constraining their dynamics to evolve within a finite domain. As a simple example of this, the top (red) curve in Fig. 1A plots the mutual information *MI*(*x*(*t*); *x*(*t* + *τ*)) for a one-dimensional Gaussian random walker as a function of time, for fixed time separation *τ* = 10 time steps. This information can be computed analytically and shown to be:

**Figure 1.**
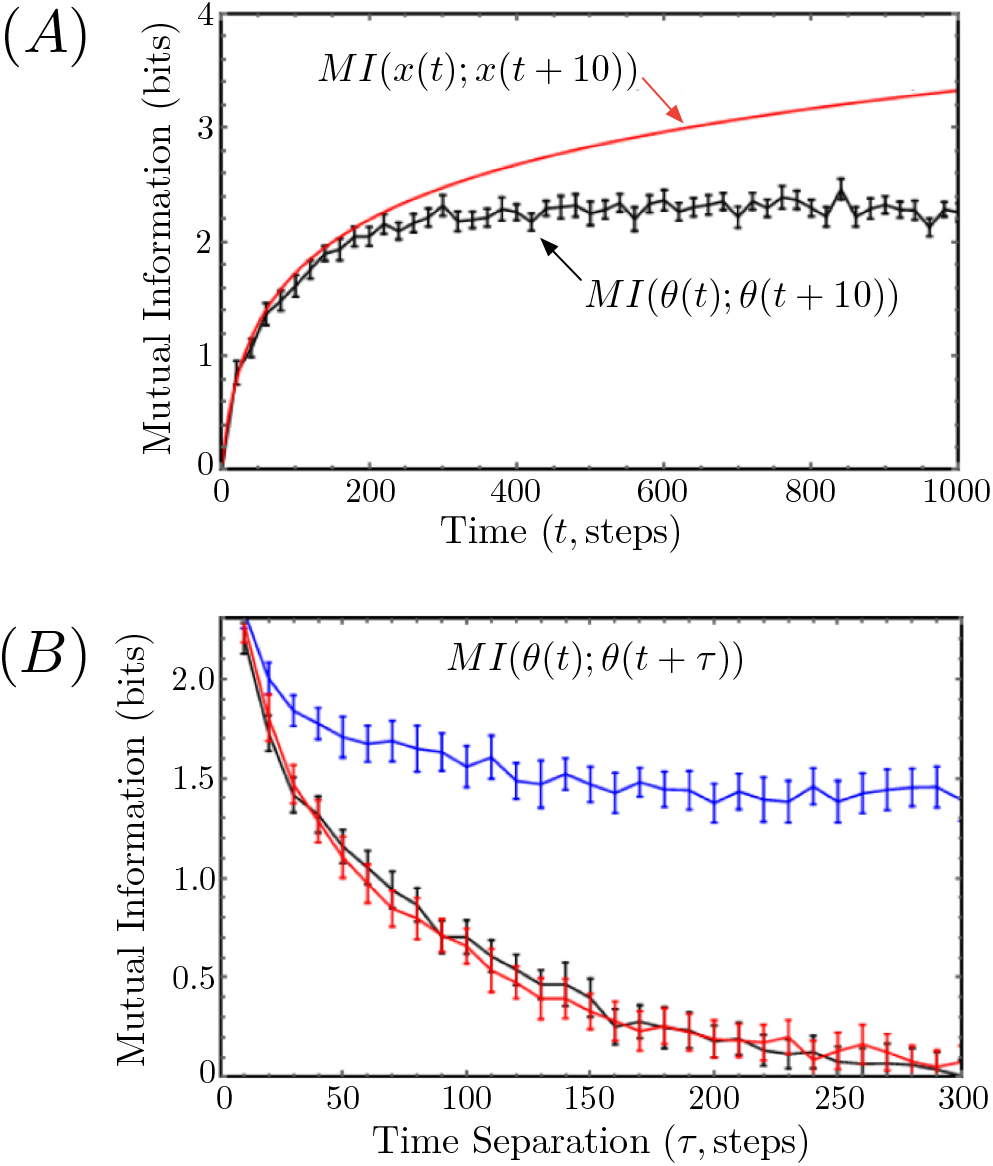
(A) The time-separated self mutual information (for fixed time separation τ = 10 time steps) of the position of a standard, one-dimensional Gaussian random walker is plotted versus the absolute time t (in red). This curve is compared with the same mutual information for a Gaussian random walker confined to the perimeter of a ring of unit radius (in black). The latter was computed numerically from an ensemble of replicate simulations using the KSG algorithm. A jackknifing procedure was used to generate multiple datasets, and the mean information is plotted with standard error bars. (Sec. 3(b) describes this procedure in more detail.) (B) The mutual information for the Gaussian random walker on a ring is plotted at steady state as a function of time separation τ. The exact information–computed from an ensemble of replicates–is plotted in black. The blue curve is computed from a single trajectory without accounting for the correlations between individually sampled pairs of time points. The red curve is computed from a single trajectory as well, but now each sampled pair of time points is at least 300 time steps apart from every other sample.

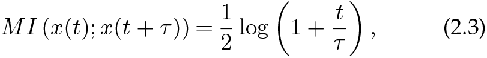

where the base of the logarithm, as usual, determines the “units” of the information (base-2 for bits, base-*e* for nats). For any fixed time separation, *τ*, this information grows logarithmically with time *t*, because the underlying distribution of *x*(*t*) has a variance that grows linearly with time as the walker tends to drift further and further from its initial position.

The bottom (black) curve in Fig. 1A plots the self mutual information *MI*(*θ*(*t*); *θ*(*t* + *τ*)) for the same Gaussian random walker when confined to the perimeter of a disk of unit radius. Now for fixed time separation, the information towards a limiting stationary value as *t → ∞*. This is because the finite circumference of the ring limits how far the walker can drift from its starting position. If the walker step size distribution has variance *σ*^2^, then the information should approach a steady state for times much longer than 2*π/σ* (in the figure, *σ*^2^ = 0.1). In other words, once the walker has had time to circle its domain many times over, the distribution of its angular position will approach stationarity.

Even if the distribution of configurations is stationary, the underlying stochastic process responsible for evolving one configuration into another will necessarily cause time-separated trajectory points to be nontrivially correlated. Fortunately, these correlations typically decay over some characteristic timescale, so if one chooses configurations separated by times much longer than that scale, they can be treated, to good approximation, as statistically independent samples of the underlying, stationary distribution.

Once again using the example of the Gaussian random walk on a ring, Fig. 1B plots the steady-state value of the mutual information *MI*(*θ*(*t*); *θ*(*t* + *τ*)) as a function of the time separation *τ*, in black. This information was computed numerically using an ensemble of replicate simulations, each of which was allowed to reach steady state before any data was collected. We compare this to the steady-state mutual information obtained from a single trajectory by naively assuming each point in the time series is a statistically independent sample (top curve in the panel, in blue). The much larger information values that result from this calculation are a result of compounding the correlations between the two time points separated by *τ* in each sample with those that arise between different sampled pairings of points. The former correlations are what we want to measure, and the latter are what we want to exclude.

Based on an estimate of the decay timescale of the ground truth mutual information in Fig. 1B, we also compute the mutual information using a subset of a single time series whose pairs of time-separated points are themselves separated from each other by a window of at least 300 time points. (For example, if one pair of points is *t*_1_ and *t*_1_ + *τ*, then the next pair, *t*_2_ and *t*_2_ + *τ*, is chosen such that *t*_2_ *− t*_1_ *≥* 300). The resulting curve, plotted in red, is statistically indistinguishable from the ground truth, as desired.

In the following sections we will continue to compare the mutual information estimated from a single time series to that estimated from an ensemble of replicate time series, so it will be useful to adopt a notation to distinguish the two. When we compute the mutual information between two random variables *X* and *Y* from a single time series, we will denote this information using the standard notation *MI*(*X*; *Y*). When we compute this same information using an ensemble of replicate time series, we will denote it *MI*(*{X}*; *{Y }*), where we have used brackets in the traditions of set theory to denote that each random variable is sampled from a set of replicate trajectories.

### (c) Golden shiner Experiments

To assess whether the salient features of the time-separated mutual information we derived for our generalized leader-follower model can emerge from an analysis of the behavior of real organisms, we recorded the motion of pairs of golden shiner fish (*Notemigonus crysoleucas*), which are well established as gregarious social organisms. The previous subsection has made it clear that our experimental methods must meet three important criteria in order for a single experimental trajectory to be suitable for an information theoretic analysis. First, the fish must be geometrically confined so that the distribution of their configurational states will tend towards a stationary distribution at long times. Second, the fish must be given sufficient time to explore their environment and reach that steady state prior to the start of the experiment. Finally, the length of the experimental data collection must be long enough **5**to ensure that there are a sufficient number of *statistically independent* datapoints to representatively sample the stationary configurational distribution of interest.

With these points in mind, we confined the fish to a roughly annular tank with an outer diameter of approximately 125.1 cm and an inner diameter of approximately 26.2 cm. In each experimental run, the two fish were given ten minutes to acclimate themselves to the tank before a high-resolution camera (mounted above) was used to record their motion for thirty minutes. A more detailed accounting of the experimental setup may be found in the appendix; at present we merely want to emphasize the aspects of the experimental design that are most pertinent to our statistical analysis.

To increase the likelihood of observing leader-follower behavior in the pairs of golden shiners, we attempted to create a more threatening set of external conditions both by increasing the intensity of the external lighting and decreasing the depth of the water in the tank. Golden shiners have been shown to prefer shaded regions, presumably to avoid predators [26, 27], and exposing the fish to conditions where predation seems more likely has been hypothesized to increase the attention each fish pays to its neighbors [26]. Changes in water depth also tend to invoke defensive behavior in social fish [28], causing them to remain in closer proximity to one another–an adaptive behavior that reduces individual risk within groups [29]. We found that pairs of shiners exposed to 230 lumens of external light intensity and a water depth of 4 cm were qualitatively much more active than fish tested under conditions of 200 lumens and a water depth of 8 cm, spending a much larger percentage of the recorded experiments circling the tank together. For future reference, we refer to the former set of experimental parameters as the agitated condition (due to the heightened anxiety level of the fish) and the latter set as the control condition.

The captured video data from the experiments was transcribed using the image-tracking software TRex, an open source platform that leverages computer vision and machine learning to identify and track moving entities [30]. With TRex, the posture of each individual fish was transcribed into a position time series featuring both the centroid and head locations of the fish in each frame of the video. The videos were recorded at 40 frames per second, so the time elapsed between most consecutive datapoints in the time series is 0.025 s, although we did have to remove small numbers of points where the image-tracking software failed to clearly recognize the individual fish and consequently did not record positional data.

To better take advantage of the radial symmetry of the tank, we re-centered the positional data to shift the coordinate origin to the middle of the tank, and then we transformed the Cartesian centroid and head locations into standard polar coordinates *r*_*i*_ and *θ*_*i*_, where *i* = 0, 1 indexes the two fish. The orientation (heading) of each fish, *Ψ*_*i*_, was computed as the angle between the positive *x*-axis and the positional vector connecting the centroid of each fish to its head. Because the tracking software sometimes had difficulty distinguishing the head of the fish from its tail, we found that the calculated heading angle would sometimes unphysically jump by nearly 180*°* between consecutive frames. To correct for this, if |*Ψ*_*i*_(*t* + 1) *− Ψ*_*i*_(*t*)| *< π −* 0.6 radians, we set *Ψ*_*i*_(*t* + 1) equal to *Ψ*_*i*_(*t*).

## 3. Results

### (a) Evidence of Follower-like Behavior

Visualization of the trajectory data revealed that, under the agitated condition, the fish transitioned synchronously in and out of two visually distinct lap-swimming behavioral modes. In the first mode, the fish swam rapid, smooth laps around the tank, with one fish apparently following closely behind the other. In the second mode, the laps were slower and subject to more irregularity in shape due to increased radial exploration, but one fish still appeared to lead the other. Trajectory segments corresponding to these two behaviors are shown in Fig. 2. Though both trajectory segments in the figure depict a single, complete lap, the smooth lap in Fig. 2A is traversed much more quickly than the irregular lap in Fig. 2B.

**Figure 2.**
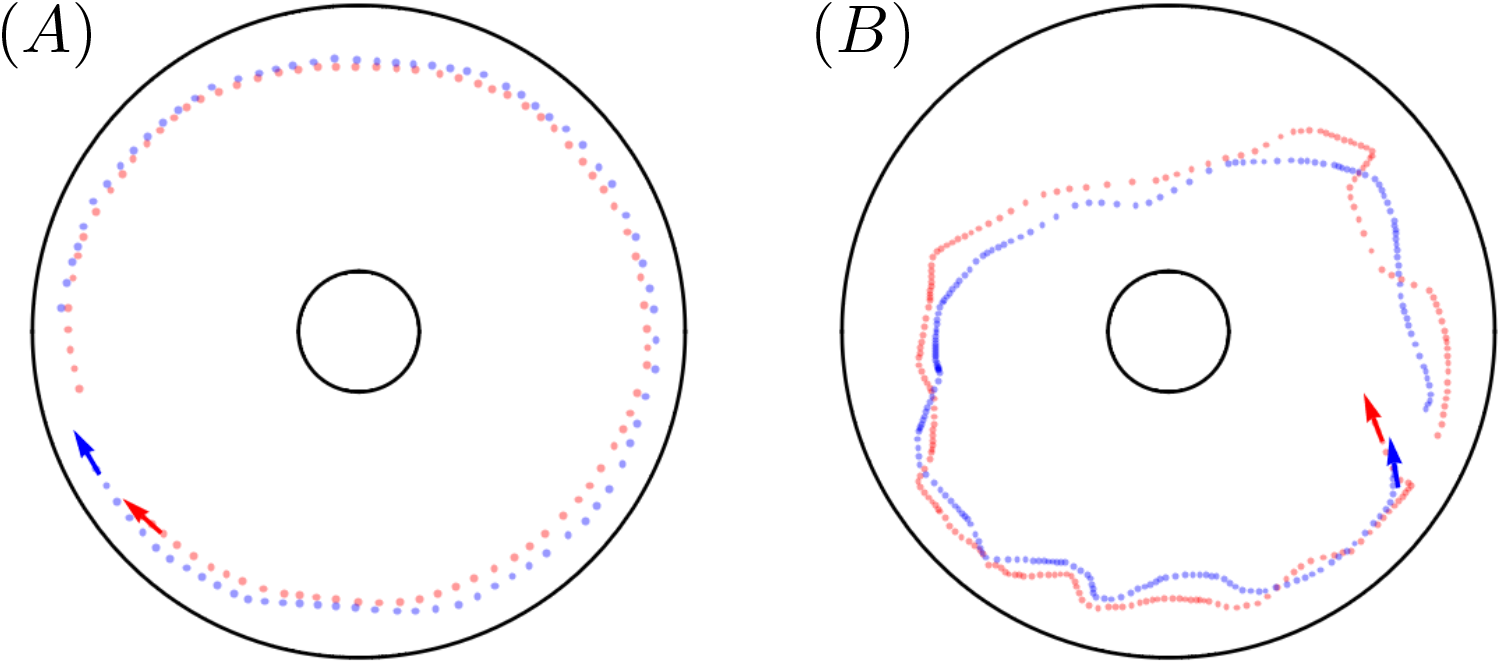
(A) A 10 s sample trajectory, taken from agitated condition experiment AS-2, in which fish-0 (red) and fish-1 (blue) swim smooth laps about the outer wall of the annular tank. (B) A 25 s sample trajectory, taken from the same experimental replicate, in which the two fish swim more erratic laps about the tank.

In addition to swapping between these two behavioral modes over the course of each experimental run, the fish also switched off between which one of them was in the lead position, further muddying the waters as to whether a true leader-follower dynamic was at play. To more clearly delineate these intervals of apparent leader-follower behavior and distinguish them from other transient behaviors, we introduce a measure of relative fish alignment, *A*_*ij*_ (*t*), that measures the angle between the heading vector of fish *i* and the vector connecting the centroid position of fish *i* to that of fish *j* at time *t*.

Defined in this manner, the alignment *A*_*ij*_ will be small (near 0) when fish *i* is swimming directly towards fish *j* and will be large (near *π*) when fish *i* is swimming directly away from fish *j*. We can use this metric to identify intervals of apparent leader-follower behavior as those where *A*_*ij*_ *≈* 0 and *A*_*ji*_ *≈ π* (or vice versa). This interpretation is illustrated schematically in the top panels of Fig. 3.

**Figure 3.**
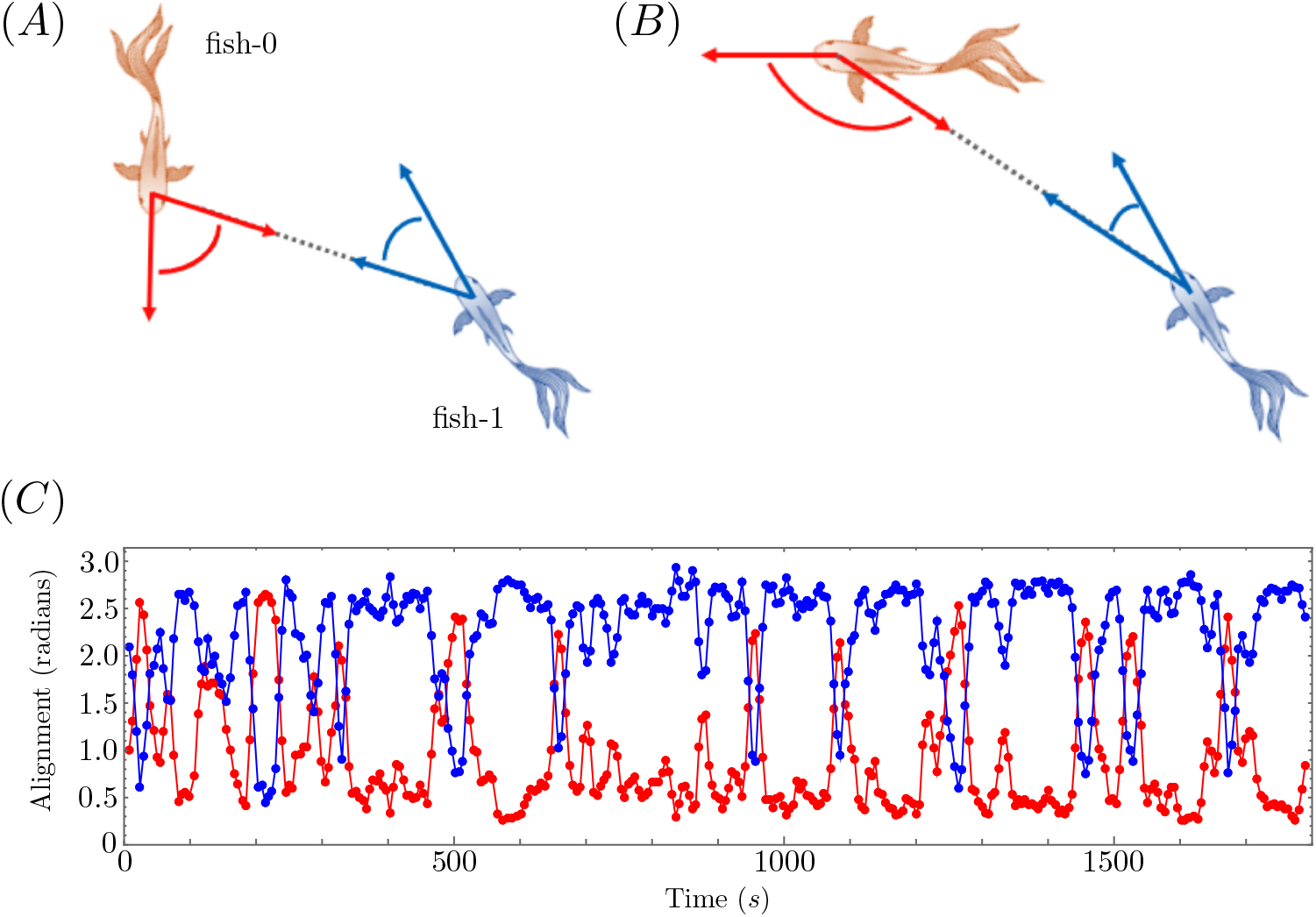
In the top-left panel, fish-0 and fish-1 are facing each other and have similar alignment angles. In the top-right panel, fish-0 leads fish-1 and fish-0 has an angle close to π where fish 1 has an angle close to 0. In the bottom panel, these alignment angles are averaged over a rolling 15 s window and are plotted for both fish over the entirety of one agitated condition experiment. Note the strong polarization in the alignments over most of the experiment and that fish-1 predominantly takes the lead position (A_10_ ≫ A_01_ over most of the time series).

In the bottom panel of Fig. 3, we plot the alignment angles of the two fish versus time for one of the agitated condition experiments. We used a rolling window averaging procedure to smooth out the noisiness of the raw data, so that each plotted point in the figure is actually the average alignment over the 15-second window centered at that time point. Over the course of this thirty minute experiment, the intervals of leader-follower-like behavior are those for which the two fish alignments polarize toward opposite extremes.

In this particular replicate, the alignment angles were strongly polarized one way or the other for nearly the entire experiment. Consequently, we shall focus on this replicate for our statistical analysis because it provides the largest quantity of useful data for interrogating whether a true leader-follower interaction can be deduced from trajectory data. Normally, cherry-picking only the best looking data is considered intellectually dishonest; but, in our case, we are merely looking for an ideal dataset for testing our statistical methods–not to draw any generalized conclusions about golden shiner interactions. For an example of more irregular golden shiner behavior, see the alignment data plotted in the appendix for a different experimental replicate.

### (b) Modeling the Data as a Simple Leader-Follower Dyad

To determine whether the observed lapping behavior of the agitated fish is driven by a genuine leader-follower dynamic, we will compare the statistics of the data to the predictions of our generalized leader-follower model from section 2(a). Since the motion of interest is approximately one-dimensional in character, we will choose 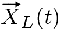 and 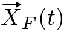 as the angular coordinates *θ*_*L*_(*t*) and *θ*_*F*_ (*t*), respectively, and assume that the two model agents are restricted to moving along a circle of fixed radius. We characterize the dynamics of the leader fish as a standard drift-diffusion process 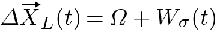, where *Ω* is a fixed angular drift term and *W*_*σ*_ (*t*) is a Gaussian white noise term with zero mean and variance *σ*^2^. The error tendency of the follower fish will similarly be modeled as Gaussian white noise, 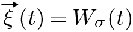. Thus, in parallel to Eq. (2.1), we define our simplified fish model with the following pair of equations:

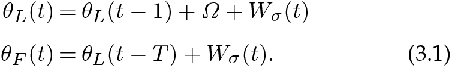

In iterating the above equations, it is to be understood that the right-hand side of both equations should always be taken modulo 2*π* to respect the periodicity of the circular system. We also assume the fish were both immobile prior to initialization, so that for *t < T*, we can set *θ*_*L*_(*t − T*) = *θ*_*L*_(0).

As before, *T* characterizes how long it takes the follower to observe the leader’s initial change in position and react by attempting to update its own position to that same location. The follower then continues to pursue the same trajectory as the leader (subject to its own errors in judgement), though it remains permanently lagged behind by *T* time steps due to the initial delayed reaction.

To account for the occasional changes in apparent leadership evident in the experiments, we add one final wrinkle to our toy model. At each time step we swap the labels on the two agents with a fixed probability *α*. Thus the fish in our model will switch between leader and follower roles on average every 1*/α* time steps. To be clear, Eq. (3.1) describes how the leader and follower positions evolve in time, but which agent obeys which equation will change back and forth during a prolonged simulation of the model. Our parameter choices for this model are derived from the experimental data of the agitated condition replicate in which the fish were observed to maintain the most robustly polarized alignment throughout the experiment (see Fig. 3). (We include the trajectory data for this replicate in the supplemental information.) If we define the angular motion of fish *i* between consecutive frames of data as *Δθ*_*i*_(*t*) *≡ θ*_*i*_(*t* + 1) *− θ*_*i*_(*t*), then we choose the angular drift of the leader, *Ω*, as the average of the random variable *Δθ*_*i*_(*t*) and the noise parameter *σ* as its standard deviation. In both cases, the average is understood to be taken for both fish over the entire experimental time series.

To give our model the same dynamical resolution as the raw video data, we define the timescales 1*/α* and *T* in units of a simulation time step of 1*/*40 s. In these units, we choose *α* = 0.00021 so that the expected number of leadership changes in thirty minutes equals the fifteen observed in the experiment, and we select *T* = 20 (0.5 s), which is just a baseline guess derived from human visual reaction measurements.

Figure 4 illustrates typical behaviors for the (ensemble-sampled) mutual information *MI*(*{θ*_1_(*t*)*}*; *{θ*_0_(*t − τ*)*}*) of our model system in three distinct cases. For each calculation, we generated 300 replicate simulations with **8***θ*_0_(0) chosen uniformly on the interval [0, 2*π*] and *θ*_1_(0) initialized at *θ*_0_(0) *−* 0.5 (modulo 2*π*, of course). Fish-0 was always initialized as the leader. We used the angular position of fish-1 at *t* = 24000 time steps, i.e., 600*s*, to ensure we were sampling from a stationary distribution in each replicate, and then, for each value of the time separation *τ*, we selected the appropriate point *θ*_0_(*t − τ*) from each replicate to build a dataset of 300 pairs of time points. To actually compute the mutual information, we jackknifed this set into ten randomly chosen subsets of 250 time point pairs and used Kraskov’s algorithm on each subset. In each of the top panels of the figure, we report the mean mutual information with standard error bars.

**Figure 4.**
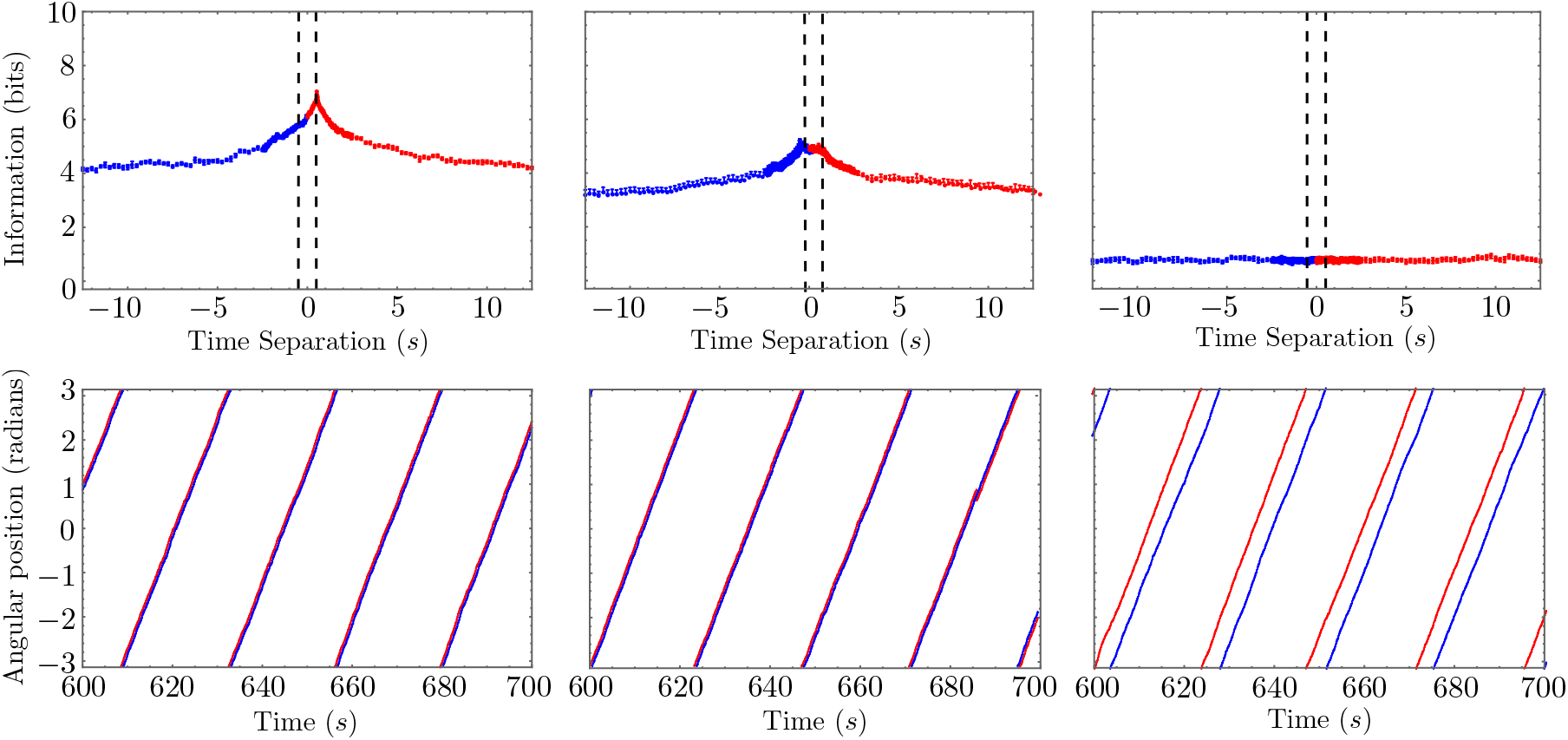
In the top three panels, we plot the time-separated mutual information MI({θ_1_(t)}; {θ_0_(t − τ)}) in our one-dimensional leader-follower model for the cases (from left to right) where there is no change in leadership (α = 0), occasional change in leadership (α = 0.00021), and where both fish behave independently (no following behavior). In all cases, we set Ω = 0067 radians per time step and σ = 0.0039 radians per time step. The bottom three panels show segments of representative trajectories corresponding to the same three cases. A dashed box highlights a change in leadership in the middle panel. Note how our mutual information metric deftly distinguishes qualitatively similar path data. (Parameters: σ = 0.156189, V = 0.2663590)

In the top-left panel, we consider the case where *α* = 0, and fish 0 acts as the leader for the entire time series. This results in a mutual information that monotonically decays away from a peak located at *τ* = 0.5*s*, corresponding exactly to the value of *T* chosen for the model. Note that the information does not appear to decay all the way to zero, but rather appears to asymptotically approach some finite, nonzero value. This is a consequence of the roughly periodic lapping behavior correlating the fish positions over very long time scales. A sufficiently large increase in the noise parameter *σ* will wash out these long-time correlations, resulting in a mutual information that fully decays towards zero.

The top-middle panel in Fig. 4 plots the same mutual information for the case where *α* = 0.00021. Now, since both fish will spend time acting as leader, we see peaks emerge at both *τ* = 0.5 and *τ* = *−*0.5 (recall that a negative time separation is equivalent to a positive time separation with the fish indices swapped). These peaks are less pronounced than in the previous case due to the stochasticity induced by the occasional leadership changes. For a sufficiently robust ensemble, the mutual information should be perfectly symmetrical about the ordinate axis since each fish has an equal opportunity to be the leader. For an ensemble with a finite number of replicates, one fish may lead a little more frequently than the other, leading to a small asymmetry. For the mutual information computed from a single trajectory time series, this asymmetry can be used to identify which fish spent more time as leader within that particular experiment.

Finally, the top-right panel in Fig. 4 considers a control case in which the fish are noninteracting (they both obey the leader dynamics of Eq. (3.1)). In this case, the mutual information is roughly constant as a function of *τ*, the small, nonzero value again resulting from the approximate periodicity of the lapping behavior. An important takeaway from this comparison is that these three qualitatively distinct mutual information patterns arise from qualitatively similar model trajectories (plotted in the bottom three panels of the figure). The first two trajectories are almost indistinguishable, except for when a leadership change occurs (highlighted by a dashed box in the figure). The third trajectory only shows a larger angular separation between the fish because the initial angular separation in their positions persists, on average, for all time. If we started the fish closer together, it would be possible to generate a trajectory that is qualitatively indistinguishable from those of the other two cases.

### (c) Estimating the Decorrelation Timescale

For the experimental fish data, we will be forced to compute the mutual information from a single time series, so we will have to employ the sort of windowing procedure outlined in Sec. 2(b). Normally, we would choose the window size, *W*, based on the decorrelation (decay) timescale of the “true” mutual information, as computed from an ensemble of replicate trajectories, but that approach obviously will not work when only a single trajectory is available. In addition, the fact that the mutual information in our model does not appear to decay to zero with time (see Fig. 4) poses another challenge to the validity of this windowing procedure. To alleviate these concerns, we propose the following procedure for computing the mutual information *MI*(*θ*_1_(*t*); *θ*_0_(*t − τ*)) from a single time series, which we will justify with our leader-follower model, where comparison with an ensemble-averaged mutual information is possible. We begin by dividing the time series for fish-1 into intervals of length *W*, for some initial choice of window size, and then we choose one time point from each interval. Rather than choose them uniformly, we treat each interval as a triangularly distributed random variable, so that we are much more likely to select time points from the middle of each interval than from the ends. Although the resulting set of time points will only be separated by *W* time steps on average when using this algorithm, we can repeat the random selection procedure to generate replicate datasets without having to reduce the size of those datasets, as we would in the jackknifing procedure discussed in the previous subsection. This is important because in many cases, experimental trajectories are woefully short, and we want to make the most of the data available.

Once we have generated our replicate datasets for *θ*_1_(*t*), we simply take the corresponding points *θ*_0_(*t − τ*) from the trajectory of fish-0 to generate our sets of paired time points and use Kraskov’s algorithm to compute the mutual information (once again reporting the average over replicate calculations with standard error bars). We then repeat the entire procedure for a larger choice of window size, and continue this search until the MI curves converge to within an acceptable tolerance.

In Fig. 5, we demonstrate this method for our simplified model in the case *α* = 0. Remarkably, even for very small window sizes of five and ten time steps (frames), we see good agreement with the ensemble-averaged result (the black curve in the figure, see also the top-left panel in Fig. 4) close to where the curve peaks. These small window sizes greatly overestimate the behavior of the information at large time separations, but we often do not care about accurately reproducing this decay. In the event that we do want to accurately reproduce the entire curve, we can simply continue to increase the window size, in which case we eventually get convergence to the ensemble-averaged result across all timescales. Note that the red curve, which agrees quantitatively with the ensemble-averaged result, corresponds to a window size of 100 time steps, or 2.5*s*, which is a reasonable estimate for the decay timescale of the black curve towards its nonzero, asymptotic limiting value.

**Figure 5.**
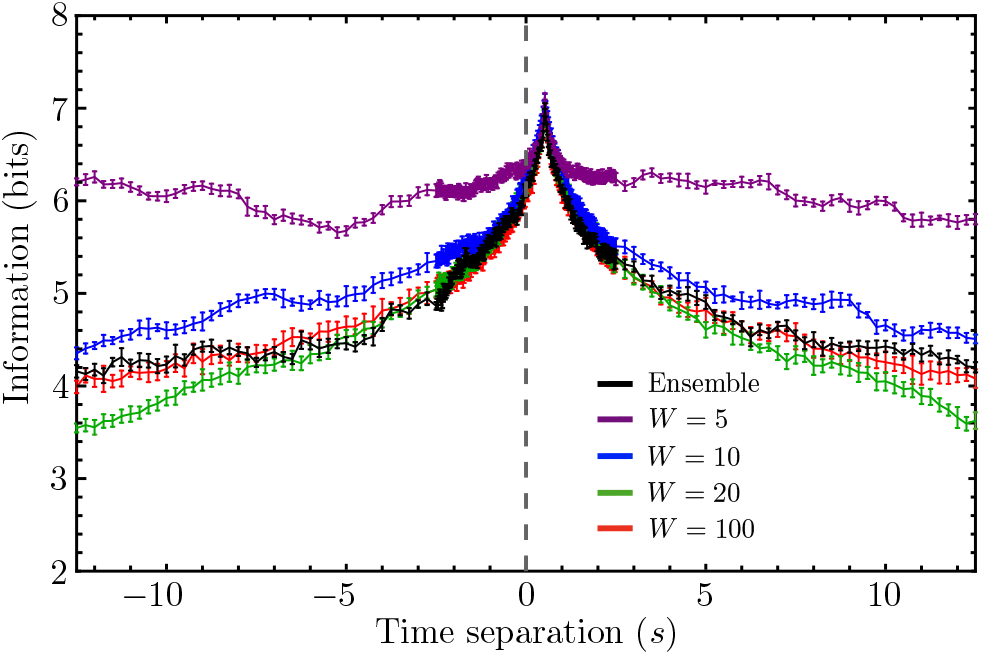
The ensemble-estimated mutual information MI({θ_1_(t)}; {θ_0_(t − τ)}) is plotted for our schematic model in black and successive single-trajectory estimates (MI(θ_1_(t); θ_0_(t − τ))) are plotted for different window sizes, ranging from W = 5 to W = 100. As expected, increasing the window between consecutively sampled pairs of time points results in convergence to the ensemble-averaged result. Even for small window sizes, however, we get surprisingly good convergence near the local maximum of the information, which is, in our case, the principal feature of interest.

### (d) Evidence of a Robust Leader-Follower Interaction

After using our phenomenological model to demonstrate how active leader-follower interactions should be expected to manifest in the mutual information shared by the time-separated angular positions of two fish (as well as how to estimate this information accurately from a single time series), we now proceed to compute the mutual information *MI*(*θ*_1_(*t*); *θ*_0_(*t − τ*)) for the time series data collected from one of our live fish experiments. As discussed previously, we selected a replicate in which the fish synchronously swam laps around the tank for most of the experiment’s duration. If the collective motion of the fish dyad in this experiment is principally driven by leader-follower-type interactions, then the mutual information computed from this dataset should most closely resemble that of our simplified model.

In Fig. 6, we plot the mutual information *MI*(*θ*_1_(*t*); *θ*_0_(*t − τ*)) for this replicate, and the resulting curve is qualitatively similar to that plotted for our schematic model in the case of no leadership changes (see the top-left panel of Fig. 4). The peak falling to the left of the ordinate axis instead of the right merely indicates that, in the experiment, fish-1 was the presumptive leader rather than fish-0, which is consistent with what we can glean from the corresponding alignment angle time series (see the bottom panel of Fig. 3).

**Figure 6.**
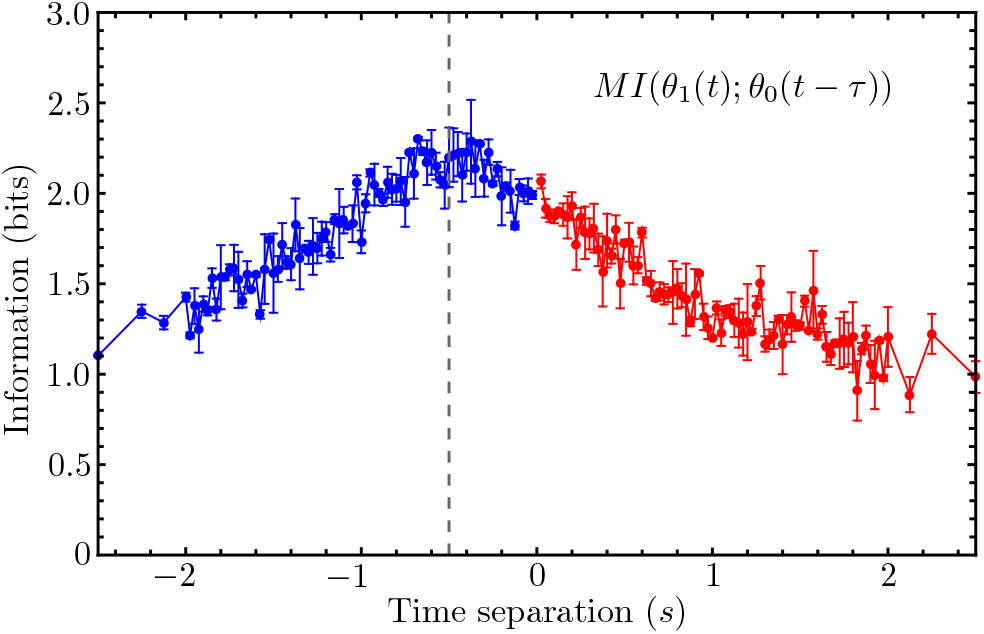
The time-separated mutual information between the angular positions of the two golden shiners, computed from a single experimental time series, is plotted for a window size W = 0.5 s. A dashed vertical line marks the estimated location of the peak. This peak occurring for τ < 0 implies that fish-1 is the leader during most of this time series.

The peak in Fig. 6 is much broader and noisier than that predicted by our model; so, to estimate the response timescale, *T*, of follower fish-0 in this experiment, it is necessary to first verify whether or not it is reasonable to treat this curve as having a single local maximum and, if it is, precisely estimate the location of this maximum.

We will accomplish this by employing the standard LOESS (locally estimated scatter plot smoothing) algorithm [31]. For each point in our mutual information plot (Fig. 6), LOESS takes the nearest *f N* datapoints, where *N* is the total number of datapoints in the plot and 0 ≤ *f* ≤ 1 is some fixed fraction of that data, and performs a locally weighted regression procedure to fit the data to a polynomial. This procedure is repeated for each datapoint in the plot, resulting in a nonparametric, systematic fitting of the overall dataset to the chosen polynomial. The important point for our purposes is that increasing the data fraction *f* will gradually smooth out more and more features of the raw data, and we want to ensure that the apparent peak in Fig. 6 is a robust enough feature to survive smoothing over a broad range of *f* values. The top panel of Fig. 7 shows the results of applying the LOESS procedure to the relevant half (*τ* ≤ 0) of the mutual information curve for three different choices of *f*. The apparent peak in the curve only begins to disappear for the largest value shown (*f* = 0.5). In the bottom panel of Fig. 7, we plot the time separation corresponding to the maximal MI value for each branch of the information curve as a function of *f*, which demonstrates that for *f* ≤ 0.5, the position of the apparent peak falls consistently within a fairly narrow band of time separation values. Averaging over these values, we extract a reaction timescale of *T ≈* 0.527 s, which is not far off from our initial model choice of *T* = 0.5 s.

**Figure 7.**
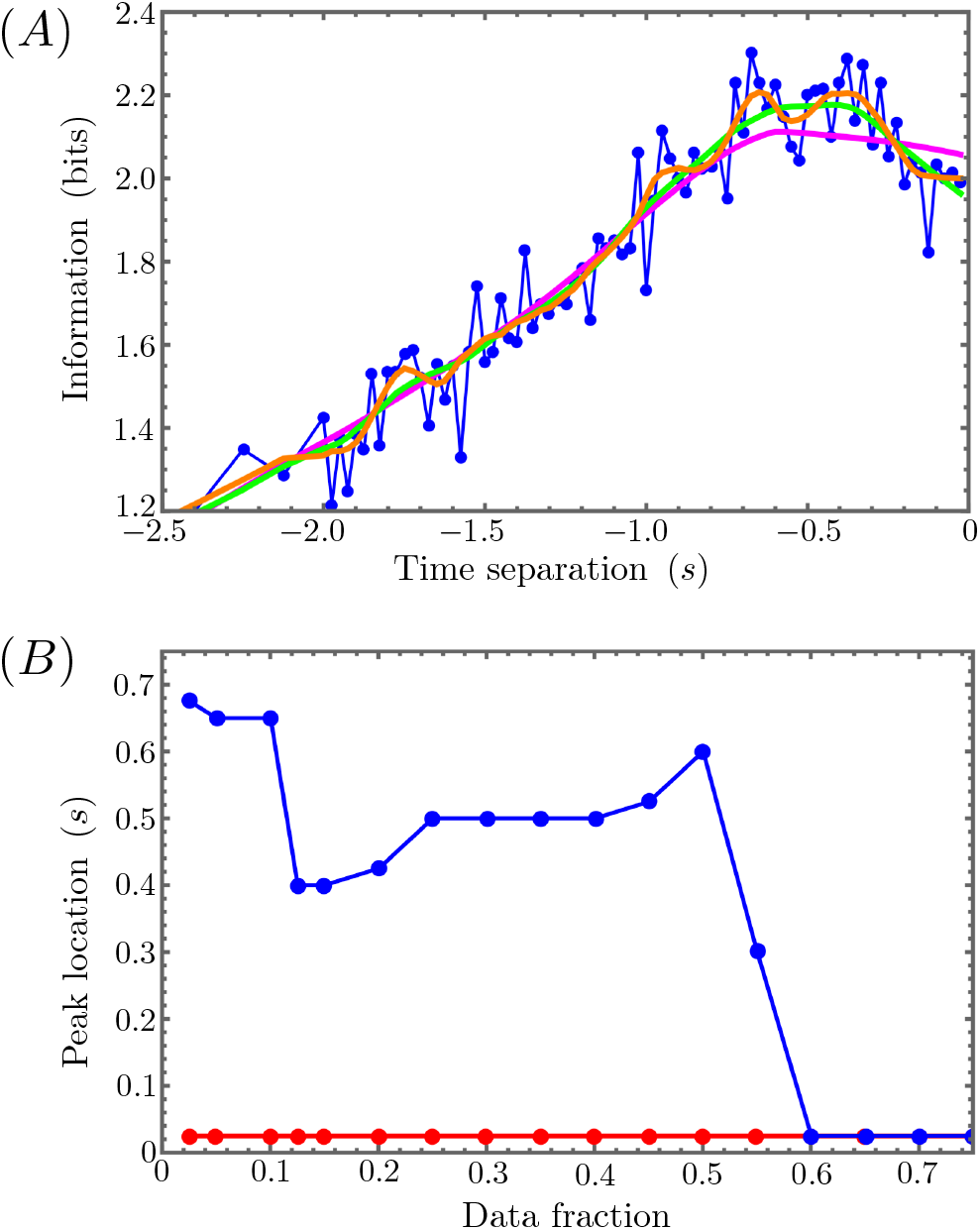
(A) LOESS smoothing of the τ ≤ 0 branch of the mutual information (in blue) plotted in Fig. 6 for data fractions f = 0.1 (orange), f = 0.2 (green), and f = 0.5 (magenta). (B) The time separation of the peak location for this same mutual information is plotted (in blue) as a function of f. For a broad range of data fractions, the peak locations falls within a narrow band of time separations. The same plot in red shows that the positive time separation branch of the MI curve in Fig. 6 has no peak for any data fraction.

## 4. Conclusions

In this paper, we have proposed a way to use mutual information, rather than transfer entropy, to quantitatively measure the extent to which asymmetric leader-follower interactions drive observed animal group behavior. Based on general considerations of what a leader-follower interaction should, at minimum, entail, we were able to argue that the time-separated mutual information between the relevant positional coordinates of a leader-follower dyad should exhibit a local maximum at some nonzero time separation corresponding to a signaling or communication timescale that is fundamental to the nature of the underlying interaction. Using a simplified, one-dimensional instantiation of our generalized model, we confirmed that this local maximum does indeed emerge at precisely the interaction timescale chosen for the model simulations.

To determine whether leader-follower interactions are present in the motion of real organisms, the experiments used to gather trajectory data must be carefully designed. To be optimal for an information theory analysis, the organisms of interest should be confined to a finite volume, and they must be permitted to explore that volume long enough to achieve a stationary distribution of organism positions. Data collection must be performed for as long a time as possible as well, since a potentially large fraction of the data will have to be discarded to ensure that each pair of time points considered can be treated as independent, uncorrelated samples.

Generally speaking, these constraints may be a tall order to implement in many systems of interest, but we have succeeded in conducting a set of experiments on pairs of golden shiners that fulfill all the desired criteria. Analyzing the trajectory data extracted from one of these experiments, we found a mutual information curve that qualitatively matched the predictions of our simple model, and from this we were able to estimate the signaling timescale of the apparent leader-follower interaction driving the synchronous lap-swimming behavior of the fish. Given the similarity of this timescale to human visual reaction times (and given the observed similarities between human and piscine vision), we can even speculate that a golden shiner tracks the motion of its confederates principally through visual cues (as opposed, for example, to sensing through the hydrodynamic fluctuations propagated by confederate motion).

## Appendix: Experimental Details

Our collaborators at the Cognitive Ecology and Behavioral Engineering Laboratory (BEL) of the U.S. Army Corp of Engineer’s Engineering Research and Development Center (ERDC) in Vicksburg, Mississippi performed the data collection and experimental management. Outside of experimentation, four hundred juvenile golden shiners were housed in an indoor holding tank with fluorescent lights on a 12h light/dark cycle. Fish were fed once daily ad libitum, after experimental individuals had been selected. To ensure a sterile environment, water quality measurements were taken daily, including temperature, dissolved oxygen, pH, conductivity, and oxidation-reduction potential (ability of water to break down contaminants). Additional weekly tests were performed to measure ammonia and nitrate content as well as carbonate hardness.

Experimental pairs were selected randomly and taken to the experimental tank in a separate temperature controlled room. An aerial view of the annular tank setup is shown in the top panel of Fig. 8. The inner diameter of the tank measured 26.2 cm and the outer diameter measured 125.1 cm. Once (unfed) experimental pairs were transferred to the tank, the fish were left undisturbed for 10 minutes to acclimate to the new environment. After the acclimation period, video data was recorded using a Basler Boost Monochrome high resolution camera for a period of 30 minutes at a frame rate of 40 frames per second and a resolution of 2912 x 2750 pixels. The still images (frames) of the video data were converted into a continuous video format using the ffmpeg library. Four replicate experiments were performed in the control setting, and five experiments were performed in the agitated setting. At the conclusion of each 30 minute experiment, fish weights and lengths were measured before euthanization.

**Figure 8.**
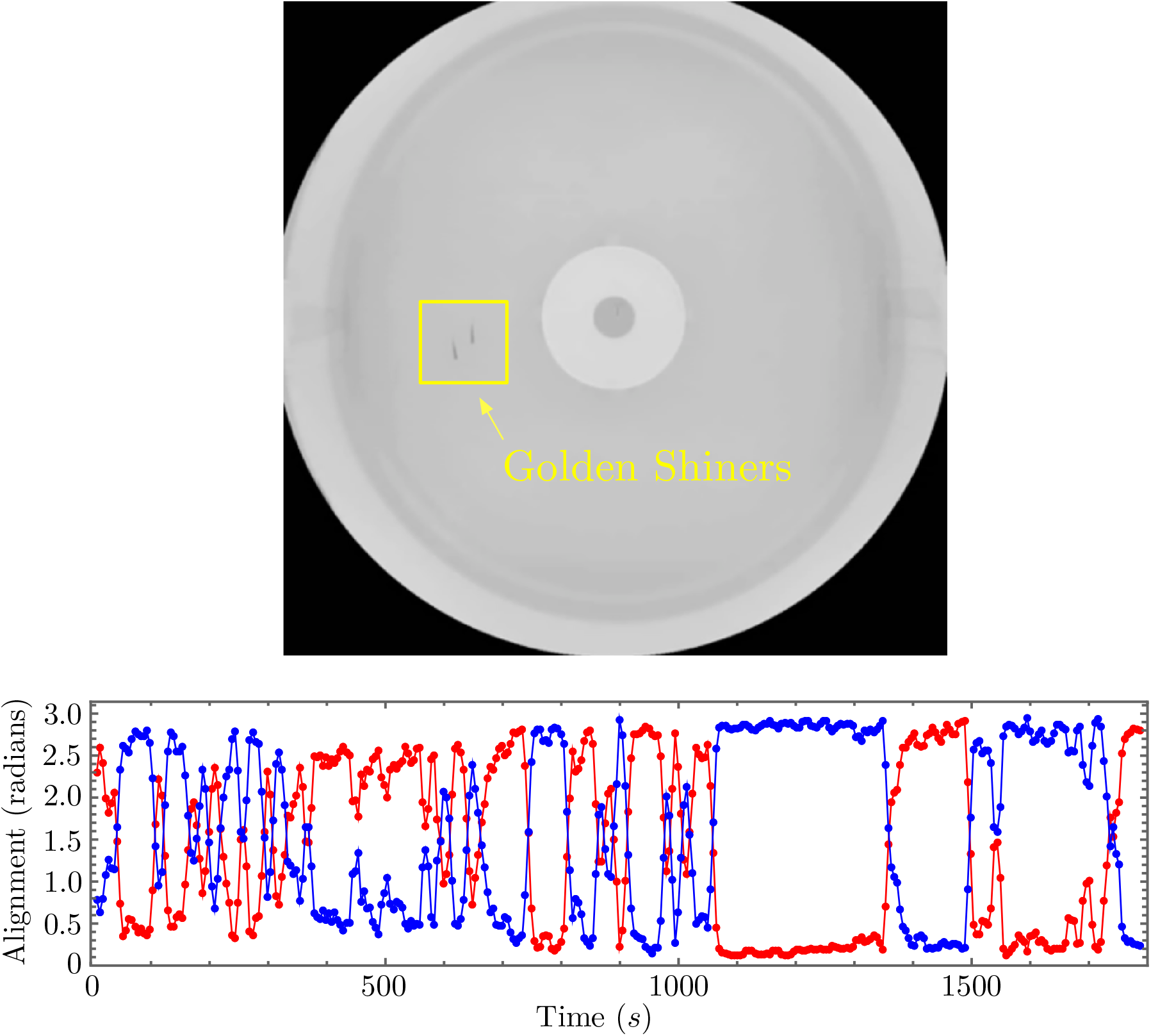
Top Panel: Raw video frame from one of the golden shiner experiments showing the annular tank and the fish dyad. Bottom Panel: A 15 s rolling window average of the alignment angles of fish-0 (in red) and fish-1 (in blue) computed over the course of a different agitated condition replicate experiment than that used throughout the rest of the paper. While the lapping behavior in this replicate is not quite as smooth and consistent as that shown in Fig. 3, long intervals of apparent leader-follower behavior can still be observed.

While we only applied our statistical analysis to the agitated replicate featuring the most consistent lapping behavior, the trajectories observed in that replicate were not behavioral outliers, as one can see from the bottom panel of Fig. 8, in which the alignment angles of the two fish are plotted for a different agitated replicate than that used for our analysis (compare this plot with that of Fig. 3).

## Ethics

The experiments performed in this study were conducted under the approval of ERDC’s Institutional Animal Care and Use Committee (IACUC). Protocol # EL-3284-2019-1.

## Data Accessibility

All the codes and algorithms used for this work may be found in the appendices of author K. Daftari’s thesis, available here: https://cdr.lib.unc.edu/concern/dissertations/8p58pq754. The experimental datasets used in the analysis are available from author K. Pilkiewicz on request.

## Authors’ Contributions

KD contributed to data curation, formal analysis, methodology, visualization, and writing - original draft; MM contributed to conceptualization, funding acquisition, methodology, visualization, and writing - review and editing; JB contributed to data curation, investigation, writing - original draft; BL contributed to conceptualization, methodology, data curation, software, and writing - review and editing. KP contributed to conceptualization, formal analysis, funding acquisition, methodology, project administration, visualization, and writing - original draft.

## Acknowledgements

The authors would like to acknowledge Dr. Elizabeth Ferguson, lead technical director of the Installations and the Operational Environment Program of the U.S. Army Corps of Engineers.

## Disclaimer

Opinions, interpretations, conclusions, and recommendations are those of the authors and are not necessarily endorsed by the U.S. Army.

